# On-site genomic epidemiological analysis of antimicrobial-resistant bacteria in Cambodia with portable laboratory equipment

**DOI:** 10.1101/2021.03.03.433720

**Authors:** Aki Hirabayashi, Hideji Yanagisawa, Hiromizu Takahashi, Koji Yahara, Philipp Boeing, Bethan Wolfenden, Vandarith Nov, Vichet Lorn, Mom Veng, Vuth Ann, Chau Darapheak, Keigo Shibayama, Masato Suzuki

## Abstract

The rapid emergence of carbapenemase-producing Gram-negative bacteria (CPGNB) is a global threat due to the high mortality of infection and limited treatment options. Although there have been many reports of CPGNB isolated from Southeast Asian countries, to date there has been no genetic analysis of CPGNB isolated from Cambodia. Sequence-based molecular epidemiological analysis enables a better understanding of the genotypic characteristics and epidemiological significance of antimicrobial-resistant (AMR) bacteria in each country, and allows countries to enact measures related to AMR issues. In this study, we performed on-site genomic epidemiological analysis of CPGNB isolated in Cambodia using a portable laboratory equipment called Bento Lab, which combines a PCR thermal cycler, microcentrifuge, gel electrophoresis apparatus, and LED transilluminator, along with the MinION nanopore sequencer. PCR targeting of major carbapenemase genes using Bento Lab revealed that two *Escherichia coli* isolates and one *Acinetobacter baumannii* isolate harbored carbapenemase genes: *bla*_NDM_, *bla*_OXA-48_, and *bla*_OXA-23_, respectively. The results of phenotypic diagnostic tests for CPGNB, such as the carbapenem inactivation method and double-disk diffusion test using a specific inhibitor of metallo-β-lactamases, were consistent with their AMR genotypes. Whole-genome sequencing analysis using MinION suggested that *bla*_NDM-5_ and *bla*_OXA-181_ genes were carried on plasmids (93.9-kb plasmid with IncFIA, IncFIB, IncFII, and IncQ1 replicons, and 51.5-kb plasmid with the IncX3 replicon, respectively) in *E. coli*, and that *bla*_OXA-23_ was encoded in two locations on the chromosome of *A. baumannii*. Plasmids carrying *bla*_NDM-5_ or *bla*_OXA-181_ in *E. coli* were highly structurally identical to plasmids prevalent in Enterobacterales in China and other countries, suggesting that they disseminated from a common evolutionary origin. Our findings demonstrate the potential impact of portable laboratory equipment on AMR bacteria research in hospitals and research centers with limited research facilities, and provide the first glimpse into the genomic epidemiology of CPGNB in Cambodia.

## Introduction

Antimicrobial-resistant (AMR) bacteria have emerged and spread all over the world. Among antimicrobials, carbapenems are one of the most reliable last-resort antimicrobials for infections caused by AMR Gram-negative bacteria. One of the mechanisms of AMR is drug inactivation mediated by acquired enzymes, such as β-lactamases [1]. Among β-lactamases, extended-spectrum β-lactamases (ESBLs) and carbapenem-hydrolyzing β-lactamase (carbapenemases) are clinically important, as ESBLs hydrolyze a broad ranges of β-lactams, including cephalosporins, and carbapenemases hydrolyze most of β-lactams, including carbapenems. β-lactamases are classified using the Ambler scheme as follows. Ambler class A includes ESBLs, such as CTX-M, as well as carbapenemases, such as KPC; class B includes metallo-β-lactamases (MBLs), such as NDM, IMP, and VIM; class C includes AmpC β-lactamases; and class D includes carbapenem-hydrolyzing oxacillinases, in which OXA-48 is prevalent in Enterobacterales, and OXA-23, OXA-24, and OXA-58 are prevalent in *Acinetobacter* spp. ESBL and carbapenemase genes are predominantly encoded on conjugative plasmids and have been transferred among Enterobacterales and other Gram-negative bacteria [2]. Moreover, carbapenemase-producing Gram-negative bacteria (CPGNB) harboring carbapenemase genes often co-harbor clinically relevant antimicrobial resistance genes, such as aminoglycoside and fluoroquinolone resistance genes [3; 4; 5]. There is great concern about the global spread of plasmids that carry multiple AMR genes and can be transferred between homogeneous and heterogeneous species.

Although CPGNB has been detected in large numbers from Southeast Asia, the publicly available information on CPGNB is limited to a small number of countries [6]. In 2015, the World Health Organization (WHO) adopted a global action plan on AMR and launched the Global Antimicrobial Resistance Surveillance System (GLASS), the first global collaborative report to standardize AMR surveillance [7]. Cambodia’s Laboratory Information System (CamLIS) was developed by the Ministry of Health in Cambodia with the support of WHO starting in 2011. As of 2018, 35 national, provincial, and referral laboratories contribute to CamLIS. Cambodia has begun its report on the GLASS report for 2020, and the actual status of AMR bacteria in the country will be revealed in the near future. To date, however, only a few studies have examined AMR bacteria clinically isolated in Cambodia, although other Southeast Asian countries are reporting increasing numbers of reports of AMR bacteria [10; 11]. So far, there has been no report on genomic epidemiology of CPGNB in Cambodia.

Here, we introduced portable laboratory equipment, Bento Lab and MinION, for on-site genomic epidemiological analysis of CPGNB in Cambodia. Bento Lab (Bento Bioworks Ltd., UK) is a DNA analysis device small enough to fit in a laptop-sized bag. It contains a PCR thermal cycler, microcentrifuge, and gel electrophoresis apparatus with LED transilluminator, and has sufficient functionality for laboratory work [15]. The MinION nanopore sequencer (Oxford Nanopore Technologies, UK) is a portable long-read sequencer with the size of a large USB memory stick. MinION was utilized for on-site genomic epidemiological analysis of the Ebola virus outbreak in West Africa in 2016 [16] and the Zika virus outbreak in the Americas in 2017 [17]. Because carbapenemase genes are mostly carried on plasmids, long-read sequencing is useful for assembling whole plasmid sequences and tracking horizontal transfer of AMR plasmids in hospitals, and local and global communities [18].

We organized an international collaborative research group from Japan, UK, and Cambodia, and successfully performed on-site genomic epidemiological analysis of CPGNB clinical isolates in locations in Cambodia where laboratory equipment is limited. Our findings demonstrate the potential impact of portable laboratory equipment on AMR bacteria research and provide the first glimpse into the genomic epidemiology of CPGNB in Cambodia.

## Materials and Methods

### Subjects and specimen collection

The outpatient clinic of National Institute of Public Health (NIPH) in Phnom Penh, Cambodia has around 10 patients in a day and 456 bacterial strains were isolated from patient specimens, such as sputum, stool, urine, pus, body fluid, and cerebral spinal fluid, in 2017. Ethical approval of this study “Genomic epidemiological analysis of AMR bacterial isolates in Cambodia” was obtained from National Ethic Committee for Health Research (NECHR), Cambodia (approval no.: 178NECHR). Two carbapenemase-producing isolates NIPH17_0020 and NIPH17_0036 of *Escherichia coli* and one carbapenemase-producing isolate NIPH17_0019 of *Acinetobacter baumannii* analyzed in this study were obtained from abdominal pus, urine, and blood of patients, respectively, at NIPH, Cambodia in 2017.

### Bacterial isolates

Bacterial species identification was performed using conventional biochemical tests and the API 20E system (bioMérieux), and antimicrobial susceptibility testing with *E. coli* ATCC 25922 as quality control was performed using BBL Sensi-Disc Susceptibility Test Discs (BD) as part of routine diagnosis at NIPH, Cambodia. Minimum inhibitory concentrations (MICs) of selected antimicrobials, including imipenem (IPM), meropenem (MEPM), ceftazidime (CAZ), cefotaxime (CTX), aztreonam (AZT), amikacin (AMK), ciprofloxacin (CPFX), and colistin (CL), against carbapenemase-producing isolates of *E. coli* (NIPH17_0020 and NIPH17_0036) and *A. baumannii* (NIPH17_0019) were further examined using the E-test strips (bioMérieux) in this study. The breakpoints for resistance (R) to antimicrobials were adopted from the Clinical and Laboratory Standards Institute (CLSI) 2020 guidelines. Carbapenemase production was examined using the carbapenem inactivation method (CIM) according to the CLSI guidelines also as routine diagnosis at NIPH, Cambodia. The double-disk diffusion tests (DDDTs) with clavulanate (CVA) and sodium mercaptoacetic acid (SMA) as specific inhibitors for extended spectrum β-lactamases (ESBLs) and metallo-β-lactamases (MBLs), respectively, were performed as previously described [19; 20; 21]. Briefly, the production of ESBLs was tested with the combination of CAZ, CTX, and amoxicillin/clavulanate (AMPC/CVA) disks (Eiken Chemical Co.), and production of MBLs was tested with the combination of IPM and SMA disks (Eiken Chemical Co.).

### PCR, whole-genome sequencing, and bioinformatics analysis

Draft genome analysis of carbapenemase-producing isolates of *E. coli* (NIPH17_0020 and NIPH17_0036) and *A. baumannii* (NIPH17_0019) using Bento Lab and MinION was performed in NIPH, Cambodia in July, 2017. Bacterial genomic DNAs (gDNAs) were extracted using the MagAttract HMW DNA Kit (Qiagen) and quantified using a Qubit 2.0 fluorometer (Thermo Fisher Scientific). PCR for selected carbapenemase genes, *bla*_NDM_ (621-bp), *bla*_KPC_ (798-bp), *bla*_IMP_ (232-bp), *bla*_VIM_ (390-bp), *bla*_OXA-48_ (438-bp), *bla*_OXA-23_ (501-bp), *bla*_OXA-24_ (246-bp), *bla*_OXA-51_ (353-bp), and *bla*_OXA-58_ (599-bp)] was performed using primers as previously described [22; 23]. Whole-genome sequencing was performed on the MinION nanopore sequencer (Oxford Nanopore Technologies) for 24 hours with the offline-capable version of MinKNOW v1.7.3 and R9.4 flow cells according to the manufacturer’s instructions.

PCR, agarose gel electrophoresis, and library preparation for MinION sequencing using the SQK-RAD002 and SQK-RAD003 kits (Oxford Nanopore Technologies) were performed using the prototype model of Bento Lab (Bento Bioworks Ltd.) consisting of a thermal cycler, microcentrifuge, and gel electrophoresis apparatus with LED transilluminator [15]. The prototype is configured slightly differently from the current commercial version, but there is no significant difference in performance (Fig. S1). Nanopore reads were basecalled using Albacore v2.1.0 (Oxford Nanopore Technologies), corrected using Genome Finishing Module v1.7 plugged-in CLC Genomics Workbench v10.1.1 (Qiagen) with default parameters of Correct PacBio Reads (beta), and assembled *de novo* using Miniasm v0.2 [24] with default parameters.

The extracted bacterial gDNAs were subsequently re-sequenced on a Illumina system in National Institute of Infectious Diseases, Japan for further error correction. Library for Illumina sequencing (insert size of 500-900 bp) was prepared using Nextera XT DNA Library Prep Kit (Illumina) and paired-end sequencing (2 x 150 bp) was performed using MiniSeq (Illumina). Illumina paired-end reads were mapped onto the on-site assembly sequences, and sequencing errors were corrected by extracting the consensus of the mapped reads five times using CLC Genomics Workbench v12.0 (Qiagen) with default parameters.

Genome sequences were annotated using the DFAST server (https://dfast.nig.ac.jp). Sequence type (ST), plasmid replicon type, and AMR genes were detected using MLST v2.0, PlasmidFinder v2.1, and ResFinder v4.1, respectively, using the CGE server (http://www.genomicepidemiology.org) with default parameters. Type IV secretion system (T4SS)-associated genes involved in conjugation were detected using TXSScan (https://galaxy.pasteur.fr/) with default parameters. Mobile gene elements (MGEs) were identified manually from CDS annotations and basically analyzed by comparing the sequences analyzed in previous studies. Linear comparisons of sequences carrying carbapenemase genes were performed using BLAST with default settings (the nucleotide collection database and the megablast program) and visualized using Easyfig v.2.2.2 (http://mjsull.github.io/Easyfig/). The annotated bacterial circular chromosomes were visualized using the CGView Server (http://cgview.ca).

Genome and plasmid sequences of carbapenemase-producing *E. coli* (NIPH17_0020 and NIPH17_0036) and *A. baumannii* (NIPH17_0019) isolated in Cambodia have been deposited at GenBank/EMBL/DDBJ under BioProject numbers PRJDB6936 and PRJDB8068.

## Results

### On-site genomic epidemiological analysis of carbapenemase-producing Gram-negative bacteria isolated in Cambodia

We stayed for five days at National Institute of Public Health (NIPH) in Phnom Penh, Cambodia in July, 2017, and set up potable laboratory equipment, including Bento Lab and MinION, in the Bacteriology laboratory with limited research facilities and no PCR machine (Fig. 1). On the first and second days, we performed the carbapenem inactivation method (CIM) test, double-disk diffusion tests (DDDTs), minimum inhibitory concentrations (MICs) measurement, PCR, and MinION sequencing on two carbapenemase-producing isolates of *E. coli* (NIPH17_0020 and NIPH17_0036) and one carbapenemase-producing isolate of *A. baumannii* (NIPH17_0019) (Fig. 1) stored in the laboratory prior to this study. On the third and fourth days, we scrutinized diagnostic testing data (Fig. 2) and analyzed sequencing data (Figs. 3-5). On the last day, we discussed the results of genotype and phenotype analysis with researchers and technicians belonging to Bacteriology laboratory of NIPH.

**Figure 1.**
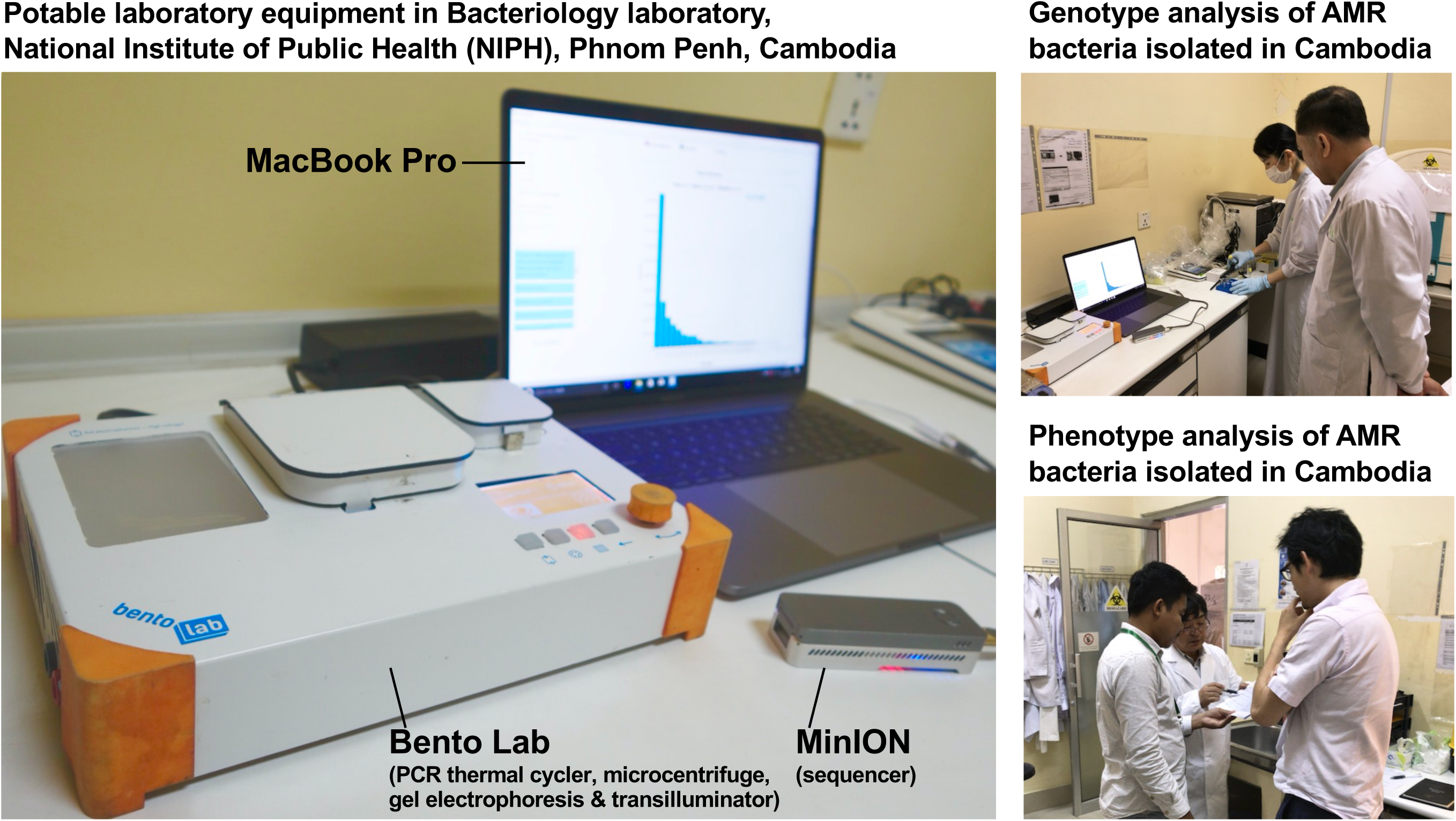
On-site genomic epidemiological analysis of AMR bacteria in Cambodia. Bento Lab and MinION were used for genotype analysis and the carbapenem inactivation method (CIM) and double-disk diffusion tests (DDDTs) were used for phenotype analysis of AMR bacteria.

**Figure 2.**
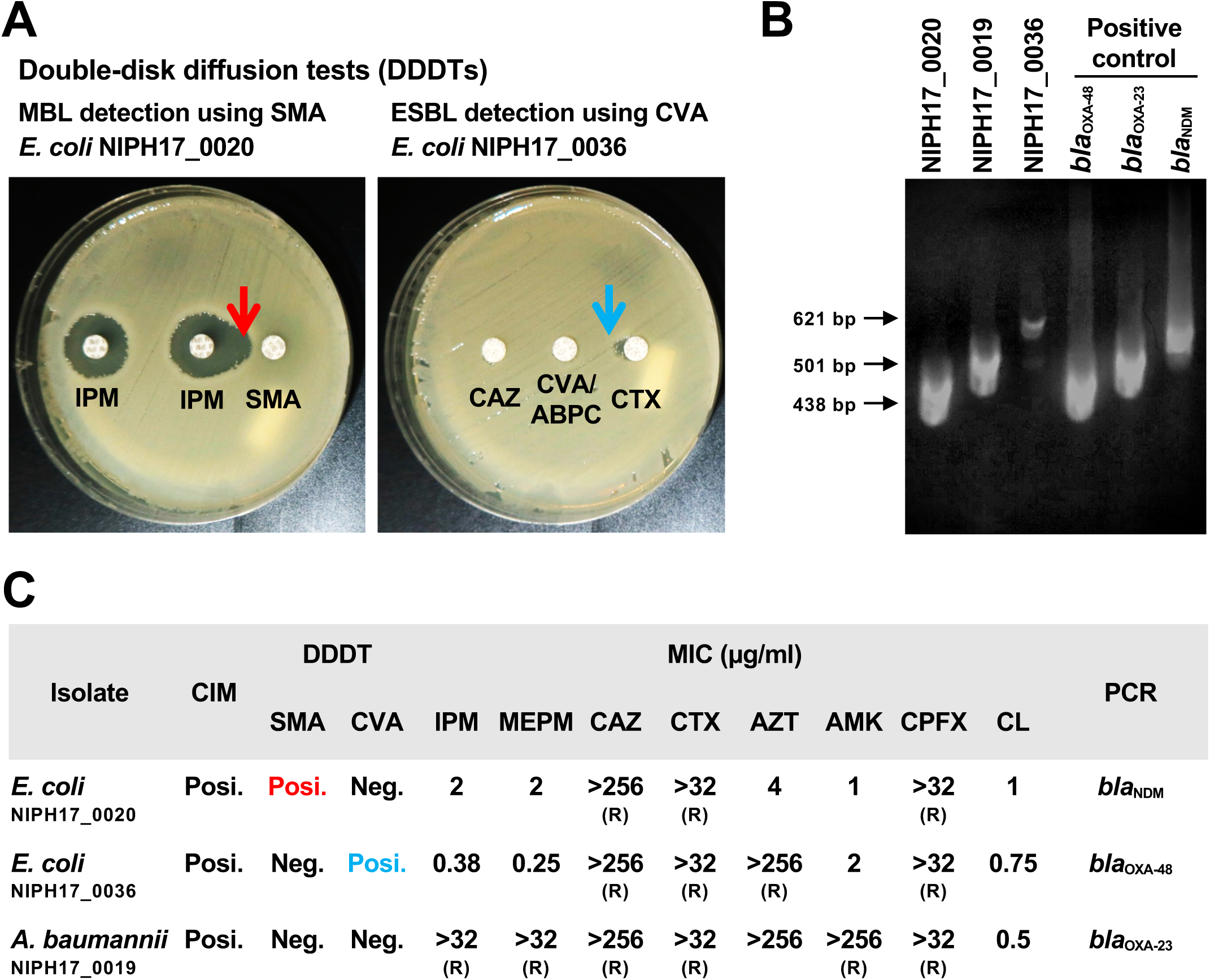
Genotype and phenotype analysis of AMR bacteria isolated in Cambodia. (A) The double-disk diffusion test (DDDT) with imipenem (IPM) and sodium mercaptoacetic acid (SMA) disks against *E. coli* NIPH17_0020 for MBL detection and DDDT with ceftazidime (CAZ), cefotaxime (CTX), and amoxicillin/clavulanate (AMPC/CVA) disks against *E. coli* NIPH17_0036 for ESBL detection. Arrows indicate β-lactamase inhibition. (B) PCR amplifications for *bla*_OXA-48_ for *E. coli* NIPH17_0036 (438 bp), *bla*_OXA-23_ for *A. baumannii* NIPH17_0019 (501 bp), *bla*_NDM_ for *E. coli* NIPH17_0020 (621 bp) and their positive controls. (C) Summary of genotype and phenotype analysis, including the carbapenem inactivation method (CIM), DDDTs with SMA or CVA, minimum inhibitory concentrations (MIC) measurement, and PCR targeting selected major carbapenemase genes. The breakpoints for resistance (R) to antimicrobials were adopted from the CLSI 2020 guidelines

The CIM test is routinely performed at NIPH, and we confirmed that all three bacterial isolates were positive for carbapenemase production. The DDDTs with sodium mercaptoacetic acid (SMA) and clavulanic acid (CVA) disks [21] showed that *E. coli* NIPH17_0020 and *E. coli* NIPH17_0036 were positive for MBL and ESBL production, respectively (Fig. 2A). PCR targeting selected major carbapenemase genes revealed that *E. coli* NIPH17_0020, *E. coli* NIPH17_0036, and *A. baumannii* NIPH17_0019 were positive for *bla*_NDM_, *bla*_OXA-48_, and *bla*_OXA-23_, respectively (Fig. 2B). The MICs of imipenem (IPM) and meropenem (MEPM) against *E. coli* NIPH17_0020, *E. coli* NIPH17_0036, and *A. baumannii* NIPH17_0019 were 2 and 2, 0.38 and 0.25, and >32 (R) and >32 µg/mL (R), respectively. Furthermore, the MICs of ceftazidime (CAZ), cefotaxime (CTX), aztreonam (AZT), amikacin (AMK), ciprofloxacin (CPFX), and colistin (CL) against *E. coli* NIPH17_0020 were >256 (R), >32 (R), 4, 1, >32 (R), and 1 µg/mL, respectively; those against *E. coli* NIPH17_0036 were >256 (R), >32 (R), >256 (R), 2, >32 (R), and 0.75 µg/mL, respectively; and those against *A. baumannii* NIPH17_0019 were >256 (R), >32 (R), >256, >256 (R), >32 (R), and 0.5 µg/mL, respectively (Fig. 2C).

Based on the on-site *de novo* assembly sequences obtained from nanopore sequencing analysis, we determined the complete structures of chromosomes and plasmids of each bacterial isolate, and detected AMR genes. As shown in Table S1, *E. coli* NIPH17_0020 had two contigs (4.78-Mb chromosome and 91.6-kb plasmid pNIPH17_0020_1); *E. coli* NIPH17_0036 had three contigs (4.68-Mb chromosome, 50.2-kb plasmid pNIPH17_0036_1, and 92.6-kb kb plasmid pNIPH17_0036_2); and *A. baumannii* NIPH17_0019 had only one contig (3.85-Mb chromosome). *E. coli* pNIPH17_0020_1 carried the *bla*_NDM-5_-like gene (96.3% identity and 3.0% gap compared with *bla*_NDM-5_: accession no. JN104597) (Fig. S2A); *E. coli* pNIPH17_0036_1 carried the *bla*_OXA-48_ family carbapenemase *bla*_OXA-181_-like gene (97.2% identity and 2.5% gap compared with *bla*_OXA-181_: accession no. CM004561) (Fig. S2B); and *A. baumannii* NIPH17_0019 harbored the *bla*_OXA-23_-like genes in two separate locations of the chromosome (97.7% identity and 2.1% gap or 93.9% and 5.6% gap compared with *bla*_OXA-23_: accession no. AY795964) (Fig. S2C).

To summarize the results, the detected carbapenemase genes had a few percent mismatch, that causes frameshift of genes, compared with their putative reference sequences (Fig. S2), thus we avoided performing CDS annotation for the on-site sequences. We at least detected AMR genes and plasmid replicons in the on-site sequences by sequence-based detection (Figs. 3–5 and Table S1). AMR genes, such as *bla*_TEM-1B_, *aadA2*, *aph(3’’)-lb*, *aac(3)-lld*, and *aph(6)-ld*, and plasmid replicons, including IncFIA, IncFIB, IncFII, and IncQ1, were detected in provisional *bla*_NDM-5_-carrying pNIPH17_0020_1 in *E. coli* NIPH17_0020 (Fig. 3), and *qnrS1* genes and IncX3 replicon were detected in provisional *bla*_OXA-181_-carrying pNIPH17_0036_1 in *E. coli* NIPH17_0036 (Fig. 4). BLAST searches of *E. coli* pNIPH17_0020_1 and *E. coli* pNIPH17_0036_1 revealed that several plasmids from Asian and Western countries were highly identical to those plasmids (Figs. 3 and 4).

**Figure 3.**
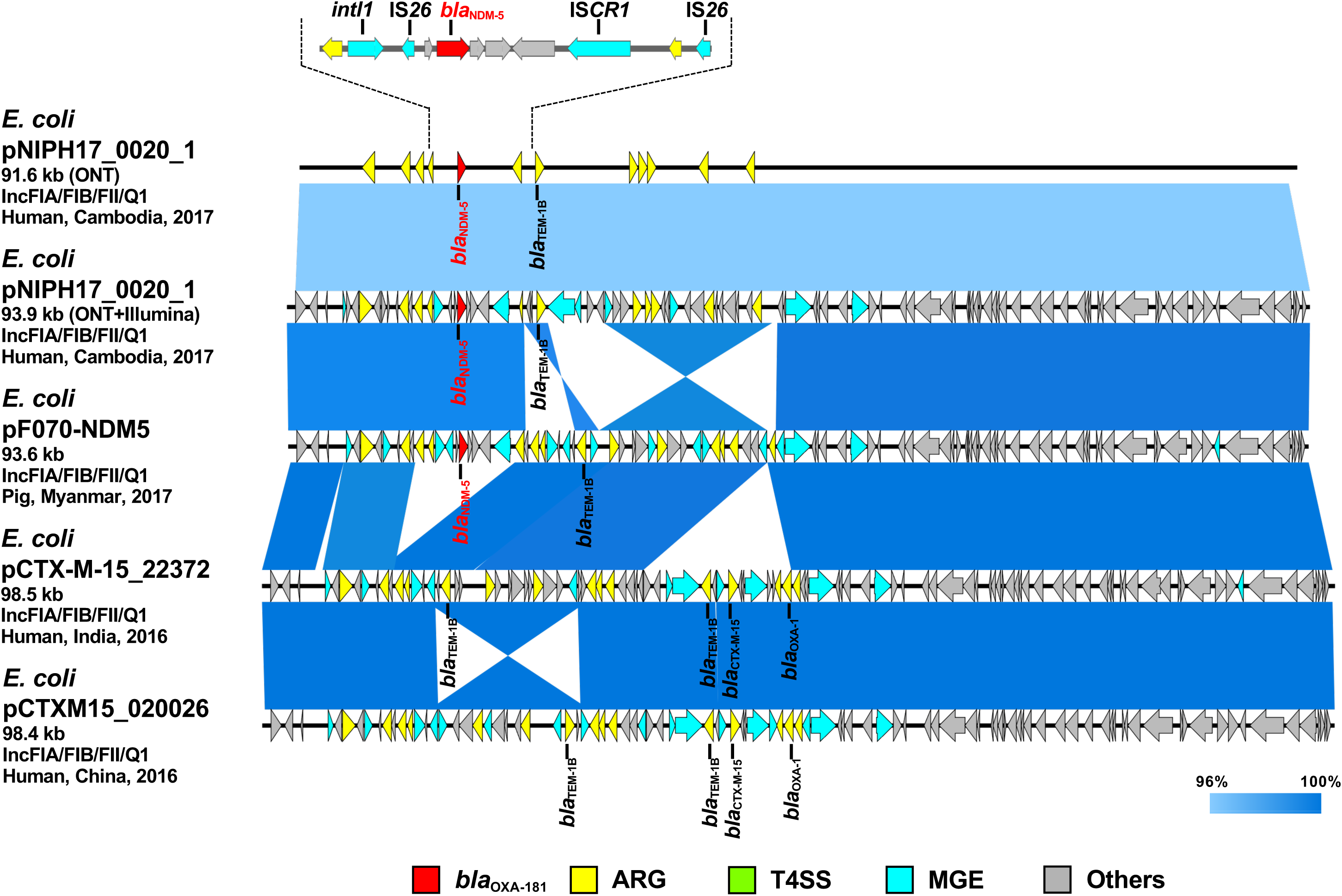
Linear comparison of related plasmid sequences from Cambodia and other countries. *E. coli* pNIPH17_0020_1 with *bla*_NDM-5_ (this study, accession no. LC483178), *E. coli* pF070-NDM5 with *bla*_NDM-5_ (accession no. AP023238), *E. coli* pCTX-M-15_22372 with no *bla*_NDM_ (accession no. CP040398), and *E. coli* pCTXM15_020026 with no *bla*_NDM_ (accession no. CP034956) are shown. Regarding *E. coli* pNIPH17_0020_1, both the uncorrected sequence (from on-site ONT analysis) and corrected sequence (from subsequent ONT+Illumina analysis), and the detailed genetic structures around *bla*_NDM-5_ are shown (upper). Red, yellow, green, blue, and gray arrows indicate carbapenemase gene (*bla*_NDM-5_), other AMR genes (ARG), type IV secretion system-associated genes involved in conjugation (T4SS), mobile gene elements (MGE), and other genes (Others), respectively. The colors in comparison of plasmids show percent identity.

**Figure 4.**
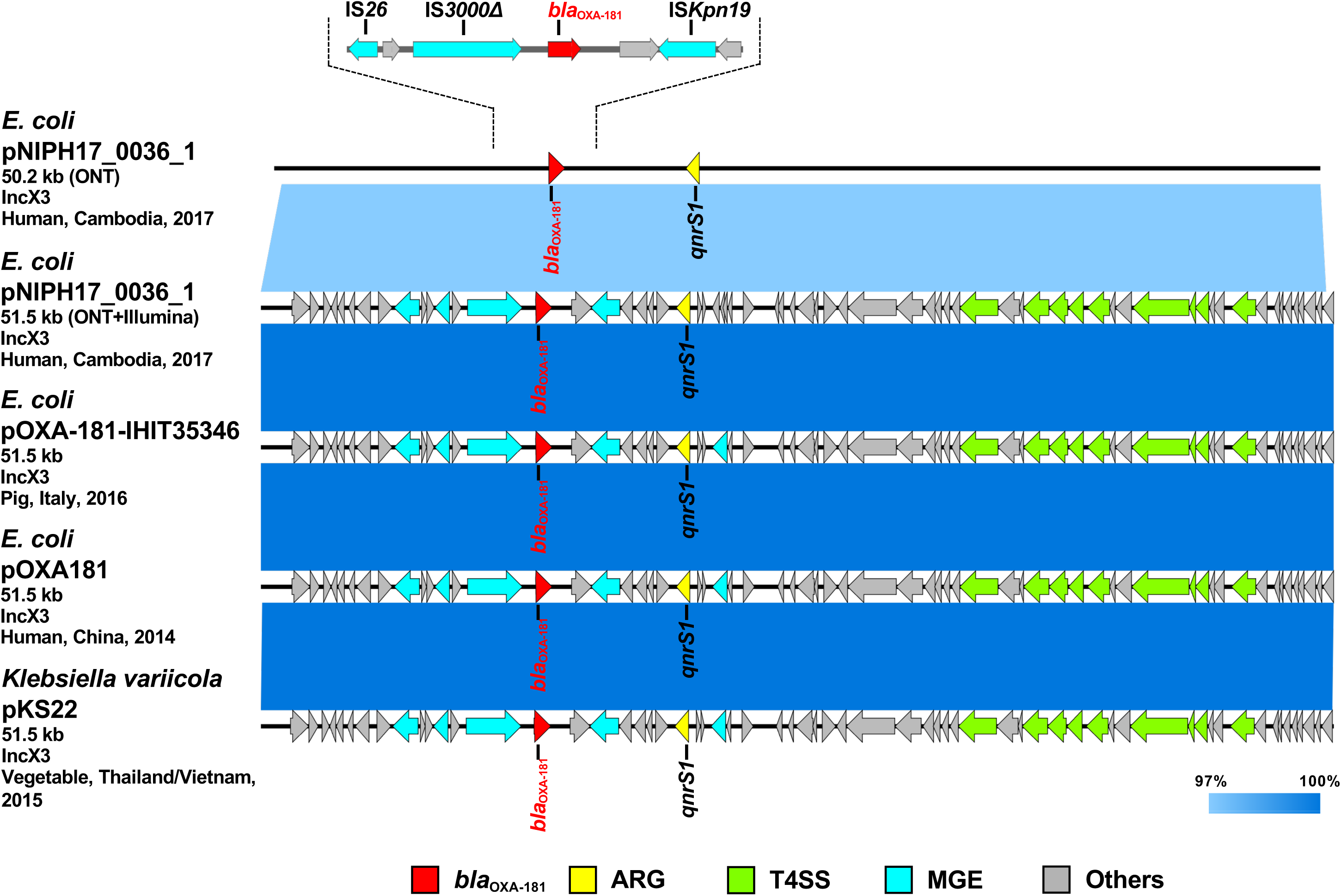
Linear comparison of related plasmid sequences from Cambodia and other countries. *E. coli* pNIPH17_0036_1 with *bla*_OXA-181_ (this study, accession no. LC483179), *E. coli* pOXA-181-IHIT35346 with *bla*_OXA-181_ (accession no. KX894452), *E. coli* pOXA-181 with *bla*_OXA-181_ (accession no. KP400525), and *Klebsiella variicola* pKS22 with *bla*_OXA-181_ (accession no. KT005457) are shown. Regarding *E. coli* pNIPH17_0036_1, both the uncorrected sequence (from on-site ONT analysis) and corrected sequence (from subsequent ONT+Illumina analysis), and the detailed genetic structures around *bla*_OXA-181_ are shown (upper). Red, yellow, green, blue, and gray arrows indicate carbapenemase gene (*bla*_OXA-181_), other AMR genes (ARG), type IV secretion system-associated genes involved in conjugation (T4SS), mobile gene elements (MGE), and other genes (Others), respectively. The colors in comparison of plasmids show percent identity.

**Figure 5.**
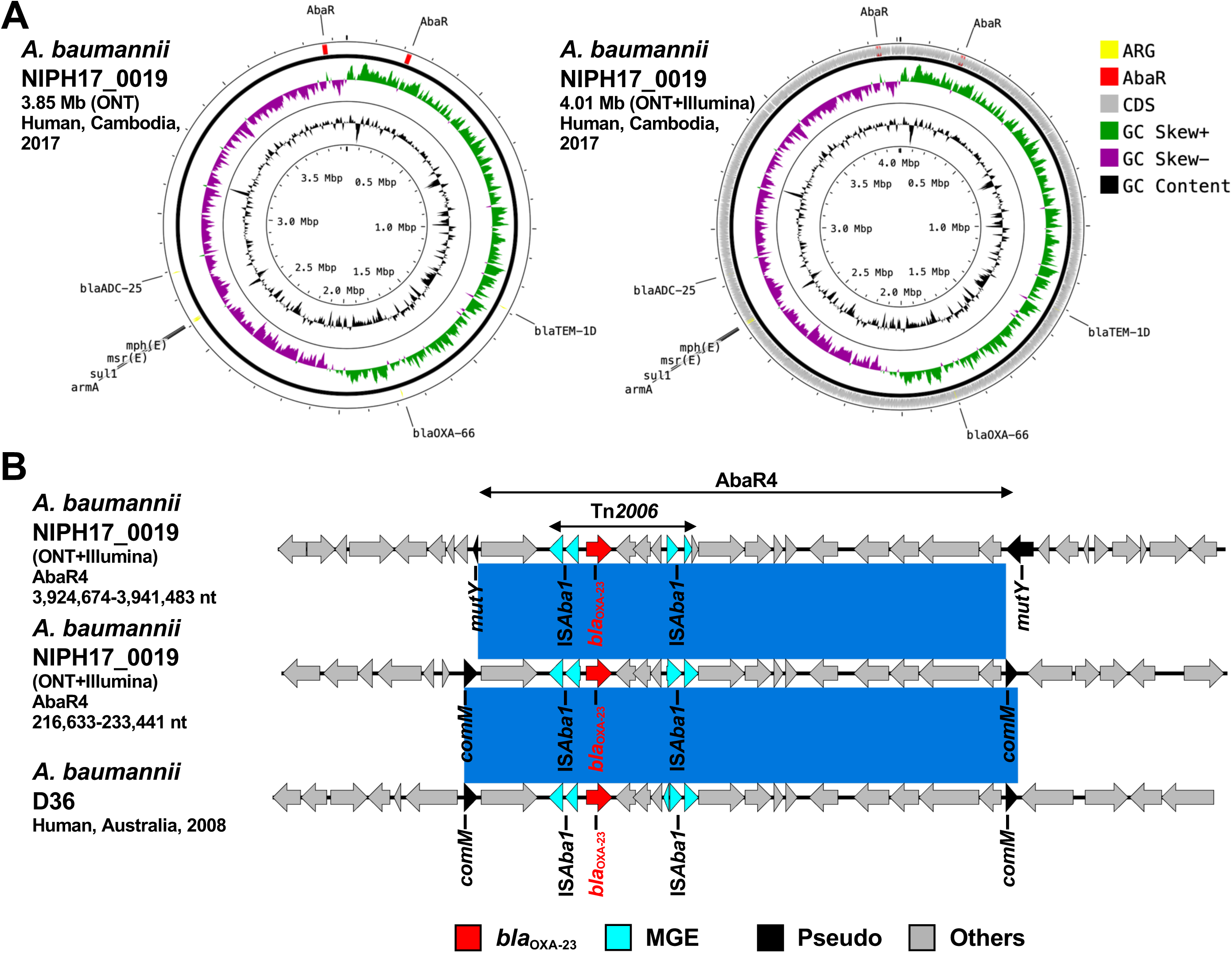
(A) Circular chromosome representation of *A. baumannii* NIPH17_0019 harboring two copies of *bla*_OXA-23_. Both the uncorrected sequence (from on-site ONT analysis) and corrected sequence (from subsequent ONT+Illumina analysis, accession no. AP024415) are shown. Yellow, red, gray, green, purple, and black indicate AMR genes (ARG), AbaR4 with *bla*_OXA-23_ (AbaR), other coding sequences (CDS), GC skew+, GC skew-, and GC content, respectively. (B) Linear comparison of AbaR4-containing genomic regions in *A. baumannii* from Cambodia and other countries. Two sets of AbaR4 with *bla*_OXA-23_ in *A. baumannii* NIPH17_0019 (3,924,674-3,941,483 nt region inserted within *mutY* and 216,633-233,441 nt region inserted within *comM* in accession no. AP024415) and AbaR4 with *bla*_OXA-23_ in *A. baumannii* D36 (41,312-58,123 nt region inserted within *comM* in accession no. JN107991) are shown. Red, blue, black, and gray arrows indicate carbapenemase gene (*bla*_OXA-23_), mobile gene elements (MGE), pseudogenes disrupted by AbaR4 insertions (Pseudo), and other genes (Others), respectively. The blue color in comparison of sequences indicates nearly 100% identity.

### Comparison of plasmids and genomic regions in carbapenemase-producing Gram-negative bacteria isolated in Cambodia with those in other countries

At a later date after the on-site analysis in Cambodia, we further performed Illumina sequencing of carbapenemase-producing isolates of *E. coli* (NIPH17_0020 and NIPH17_0036) and *A. baumannii* (NIPH17_0019), corrected the on-site *de novo* assembly sequences using Illumina reads, and compared the on-site and error-corrected sequences (Figs. 3, 4, 5A and S3). The on-site sequences of *E. coli* pNIPH17_0020_1 (provisional 91.6-kb *bla*_NDM-5_-carrying plasmid with IncFIA/FIB/FII/Q1 replicons) and *E. coli* pNIPH17_0036_1 (provisional 50.2-kb *bla*_OXA-181_-carrying plasmid with IncX3 replicons) were highly identical with their error-corrected sequences (96.48% identity in 100% region of the error-corrected 93.9-kb pNIPH17_0020_1: accession no. LC483178 and 96.70% identity in 100% region of the error-corrected 51.5-kb pNIPH17_0036_1: accession no. LC483179, respectively) (Figs. 3 and 4). Moreover, the on-site sequences of *bla*_OXA-23_-containning chromosomal regions of *A. baumannii* NIPH17_0019 were also highly identical with their error-corrected sequences (99.88% identity in 100% region of 216,633-233,441 nt and 99.84% identity in 100% region of 3,924,674-3,941,483 nt in their error-corrected chromosome of NIPH17_0019: accession no. AP024415) (Figs. S3A and S3B). Although there were a few percent differences between the on-site and error-corrected sequences, the best match types of AMR genes and plasmid replicons detected from the reference libraries were consistent, respectively (Figs. 3, 4 and 5A).

*E. coli* NIPH17_0020 belonged to sequence type 410 (ST410) in multilocus sequence typing (MLST) analysis and had one 93.9-kb plasmid pNIPH17_0020_1 with a backbone comprising IncFIA/FIB/FII/Q1 that carried multiple AMR genes, such as β-lactamase (*bla*_NDM-5_ and *bla*_TEM-1B_) and aminoglycoside resistance genes [*aac(3)-lld*-like, *aadA2, aph(3”)-lb, aph(6)-ld*] (Table S1). *E. coli* pNIPH17_0020_1 was structurally highly identical with plasmid pPF070-NDM5 (accession no. AP023238) in *E. coli* isolated from a pig in Myanmar in 2017, plasmid pCTX-M-15_22372 (accession no. CP040398) in *E. coli* isolated from a human in India in 2016, and plasmid pCTXM15_020026 (accession no. CP034956) in *E. coli* from a human in China in 2016 (99.8% identity in 92-94% regions of pNIPH17_0020_1) (Fig. 3). pNIPH17_0020_1 contained several mobile gene elements (MGEs), including IS*26* and IS*CR1,* surrounding *bla*_NDM-5_ (Fig. 3 upper). pF070-NDM5 carried *bla*_NDM-5_ with the same MGEs, whereas pCTX-M-15_22372 and pCTXM15_020026 did not carry *bla*_NDM-5_ (Fig. 3).

*E. coli* NIPH17_0036 also belonged to ST410 in MLST analysis and had two plasmids; one of them, 51.5-kb IncX3 plasmid pNIPH17_0036_1 carrying AMR genes, including *bla*_OXA-181_ and *qnrS1* (Table S1), was structurally nearly identical with plasmid pOXA-181-IHIT35346 (accession no. KX894452) in *E. coli* isolated from a pig in Italy in 2016 and plasmid pOXA181 (accession no. KP400525) in *E. coli* isolated from a human in China in 2014 [25], as well as plasmid pKS22 (accession no. KT005457) in *Klebsiella variicola* isolated from a fresh vegetable imported from Thailand or Vietnam (99.9% identity in 100% regions of pNIPH17_0036_1) (Fig. 4). *bla*_OXA-181_ in pNIPH17_0036_1 was surrounded with several MGEs, including IS*Kpn19*, IS*3000*, and two copies of IS*26* (Fig. 4 upper). *E. coli* NIPH17_0036 had another plasmid pNIPH17_0036_2 (94.8-kb IncFIA/FIB/FII/Q1 plasmid, accession no. LC603215) carrying *bla*_CTX-M-15_, *bla*_OXA-1_, *bla*_TEM-1B_, other β-lactamase genes, and multiple aminoglycoside resistance genes (Table S1).

*A. baumannii* NIPH17_0019 belonged to ST571 in MLST analysis and harbored the *bla*_OXA-23_ genes in two separate regions on the chromosome (accession no. AP024415) (Fig. 5A). Both copies of *bla*_OXA-23_ were located in Tn*2006* in AbaR4. AbaR is the resistance island mediating AMR genes in *A. baumannii* [36]. Comparison of the genetic environment around AbaR4 was performed between *A. baumannii* NIPH17_0019 and AbaR4-harboring *A. baumannii* D36 (accession no. JN107991), which was isolated from a human in 2008 in Australia (Fig. 5B). The result revealed that two AbaR4 in *A. baumannii* NIPH17_0019 were highly identical to that of *A. baumannii* D36 (99.84-99.88% identity in 100% regions) (Fig. 5B). Interestingly, AbaR4 was integrated into the *comM* gene in *A. baumannii* D36, whereas AbaR4 were integrated into the *comM* gene (for AbaR4 of 216,633-233,441 nt region) and *mutY* gene (for AbaR4 of 3,924,674-3,941,483 nt region) in *A. baumannii* NIPH17_0019.

## Discussion

In this study, we successfully conducted genomic analysis of carbapenemase-producing Gram-negative bacteria (CPGNB) in Cambodia and showed the first glimpse into the genomic epidemiology of CPGNB in Cambodia. The main analysis was performed on site using portable laboratory equipment: Bento Lab and MinION. The nearly complete genomes, including plasmids, of carbapenemase-producing *E. coli* and *A. baumannii* isolates were determined using the MinION nanopore sequencing data, and detection of AMR genes and plasmid replicons, and sequence similarity search in the public database were performed at the laboratory with limited research facilities in Cambodia. Since the nanopore sequencing technology is still evolving and the accuracy of sequencing is not perfect at present due to the a few percent rates of indels and substitution [26], it was necessary to combine other sequencing technologies, such as Illumina sequencing by synthesis, for further molecular typing analysis of bacteria that requires more accurate sequences at the single nucleotide level.

The carbapenem inactivation method (CIM) and double-disk diffusion test (DDDT) are simple, inexpensive, and useful, especially in developing countries where materials and facilities are limited. In this study, the results of both tests were reasonable, as validated by subsequent genetic analysis (Fig. 2). The CIM test is routinely performed in NIPH, Cambodia, and NIPH had detected and stored carbapenemase-producing bacterial isolates prior to this study. The DDDTs with SMA and CVA disks clearly detected production of MBL in *E. coli* NIPH17_0020 and ESBL in *E. coli* NIPH17_0036, respectively (Fig. 2A). The CIM and DDDT are important for screening for CPGNB because MICs of carbapenems against CPGNB are not always high, as bacteria with carbapenem-hydrolyzing oxacillinase genes, such as *bla*_OXA-48_ or its variants. The MICs of carbapenems against *E. coli* NIPH17_0036, which harbored *bla*_OXA-181_ on its plasmid (Fig. 4), were relatively low, whereas CIM yielded a positive result (Fig. 2C). OXA-48 is capable of weakly hydrolyzing carbapenems while maintaining its activity against broad-spectrum cephalosporins. The *bla*_OXA-181_ gene is a variant of *bla*_OXA-48_, and the hydrolysis activity of OXA-181 for β-lactams are similar to that of OXA-48 [37].

We sequenced carbapenemase-producing *E. coli* and *A. baumannii* isolates on site using MinION and Bento Lab, and revealed that *E. coli* NIPH17_0020 and *E. coli* NIPH17_0036 harbored *bla*_NDM-5_ and *bla*_OXA-181_ on their plasmids pNIPH17_0020_1 and pNIPH17_0036_1, respectively (Figs. 3 and 4) and that *A. baumannii* NIPH17_0019 harbored two copies of *bla*_OXA-23_ on its chromosome (Fig. 5). While the MinION control software, MinKNOW requires a constant internet connection, we were provided with the offline-capable version of MinKNOW from the company and used on-site analysis. For *de novo* assembly using nanopore long-read data, we used the Miniasm software, which is fast and computationally inexpensive [24]. Since Minasm assembles without error correction, MinION reads were error corrected using the pipeline of CLC Genomics Workbench for long-read sequencers prior to *de novo* assembly. Although the resulted assembly sequences obtained from the on-site analysis still contained a few percent of errors, it was sufficient for subsequent molecular epidemiological analysis. Although the resulted assembly sequences obtained from the on-site analysis contained a few percent of errors, it was sufficient for subsequent molecular epidemiological analysis, and the error-corrected sequences using Illumina sequencing were subsequently used to confirm the results of the on-site analysis (Figs. 3–5).

*E. coli* pNIPH17_0020_1 [93.9-kb IncFIA/FIB/FII/Q1 plasmid with *bla*_NDM-5_, accession no. LC483179] was structurally highly identical to *E. coli* pPF070-NDM5 in Myanmar (accession no. AP023238) (Fig. 3), and harbored several mobile gene elements (MGEs), including IS*26* and IS*CR1* surrounding *bla*_NDM-5_ (Fig. 3 upper). The *bla*_NDM-5_-conaining regions between IS*26* and IS*CR1* in pNIPH17_0020_1 and pPF070-NDM5 were identical with those of IncFII plasmids, such as *E. coli* pM109_FII in Myanmar (accession no. AP018139), *E. coli* pGUE-NDM in France (accession no. JQ364967), and *K. pneumoniae* pCC1409-1 in South Korea (accession no. KT725789), implying that *bla*_NDM-5_ was deseminated via plasmids and also MGEs, such IS*26*, among Enterobacterales over the world [27].

*E. coli* pNIPH17_0036_1 [51.5-kb IncX3 plasmid with *bla*_OXA-181_, accession no. LC483179] was structurally nearly identical with *E. coli* pOXA-181-IHIT35346 in Italy (accession no. KX894452), *E. coli* pOXA181 in China (accession no. KP400525), and *K. variicola* pKS22 in Thailand/Vietnam (accession no. KT005457) (Fig. 4). Our analysis revealed that *bla*_OXA-181_-carrying IncX3 plasmids widespread among Enterobacterales worldwide were also present in Cambodia. pKS22 was detected in coriander imported from Thailand or Vietnam, and the international fresh vegetable trade is suspected to be a route for the spread of AMR bacteria [28]. Since Cambodia is geographically and culturally close to Thailand and Vietnam, there is a possibility of transmission of AMR bacteria through foods. However, this study was very small, so further analysis with larger numbers of bacterial isolates in Cambodia according to One-Health approaches will be necessary to characterize AMR bacteria in this country.

*A. baumannii* NIPH17_0019 belonging to ST471 harbored two copies of *bla*_OXA-23_ in the chromosome (Fig. 5A). According to a previous study of carbapenem-resistant *A. baumannii* [14], ST2 is the most prevalent genotype and *bla*_OXA-23_ is the most widespread carbapenem resistance gene in the world. ST471 belongs to clonal complex 2, and there was a report that ST471 strains harboring *bla*_OXA-23_ were widespread in medical settings in Vietnam [29]. Generally, carbapenem-hydrolyzing oxacillinases hydrolyze carbapenems weakly and do not contribute to strong carbapenem resistance on their own. However, the elevated expression of the oxacillinase genes by the upstream insertion of IS, such as ISAba1 in Acinetobacter spp., which serves as a promoter for the downstream genes and leads to strong resistance [30; 31]. Two copies of *bla*_OXA-23_ in *A. baumannii* NIPH17_0019 were located downstream of IS*Aba1*, respectively (Fig. 5B). Both *bla*_oxa-23_ were included in Tn*2006*, which is commonly found 4.8-kb transposon in *Acinetobacter* app., with a central segment of 2,445-bp flanked by two reverse-oriented copies of IS*Aba1* [32]. Tn*2006*-cantaining AbaR4 is frequently inserted within the *comM* gene in the chromosome of *A. baumannii* international clones [33; 34; 35], and the *comM* gene is known as the most-preferred hotspot for AbaR insertions [36]. One of the insertion sites of AbaR4 (3,924,674-3,941,483 nt region) in *A. baumannii* NIPH17_0019 was not the *comM* gene but the *mutY* gene, adenine DNA glycosylase (Fig. 5B). According to a previous study [36], AbaR insertions at *acoA*, *pho,* and *uup* genes were occasionally observed; however, *mutY* has not been previously reported as the insertion site.

## Conclusion

Based on our on-site genomic epidemiological analysis of carbapenemase-producing Gram-negative bacteria in Cambodia, we revealed for the first time that plasmids and MGEs carrying clinical relevant carbapenemase genes reported in other countries have also been spreading in Cambodia. Bento Lab and MinION are useful for genomic analysis and surveillance of AMR bacteria in hospitals and research centers with limited facilities.

## Acknowledgements

This work was supported by Grants-in-Aid for the Research Program on Emerging and Re-emerging Infectious Diseases (JP20fk0108093, JP20fk0108139, JP20fk0108133, and JP20wm0225008 for MS, and JP20fk0108061 for KS) and the Japan Initiative for Global Research Network on Infectious Diseases (J-GRID) (JP18fm0108003 and JP19fm0108006 for KS) from the Japan Agency for Medical Research and Development (AMED). This work was also supported by the Cooperative Research Grant from Research Center on Tropical Disease, Institute of Tropical Medicine, Nagasaki University (2019-Ippan-20).

---

**Table S1.**
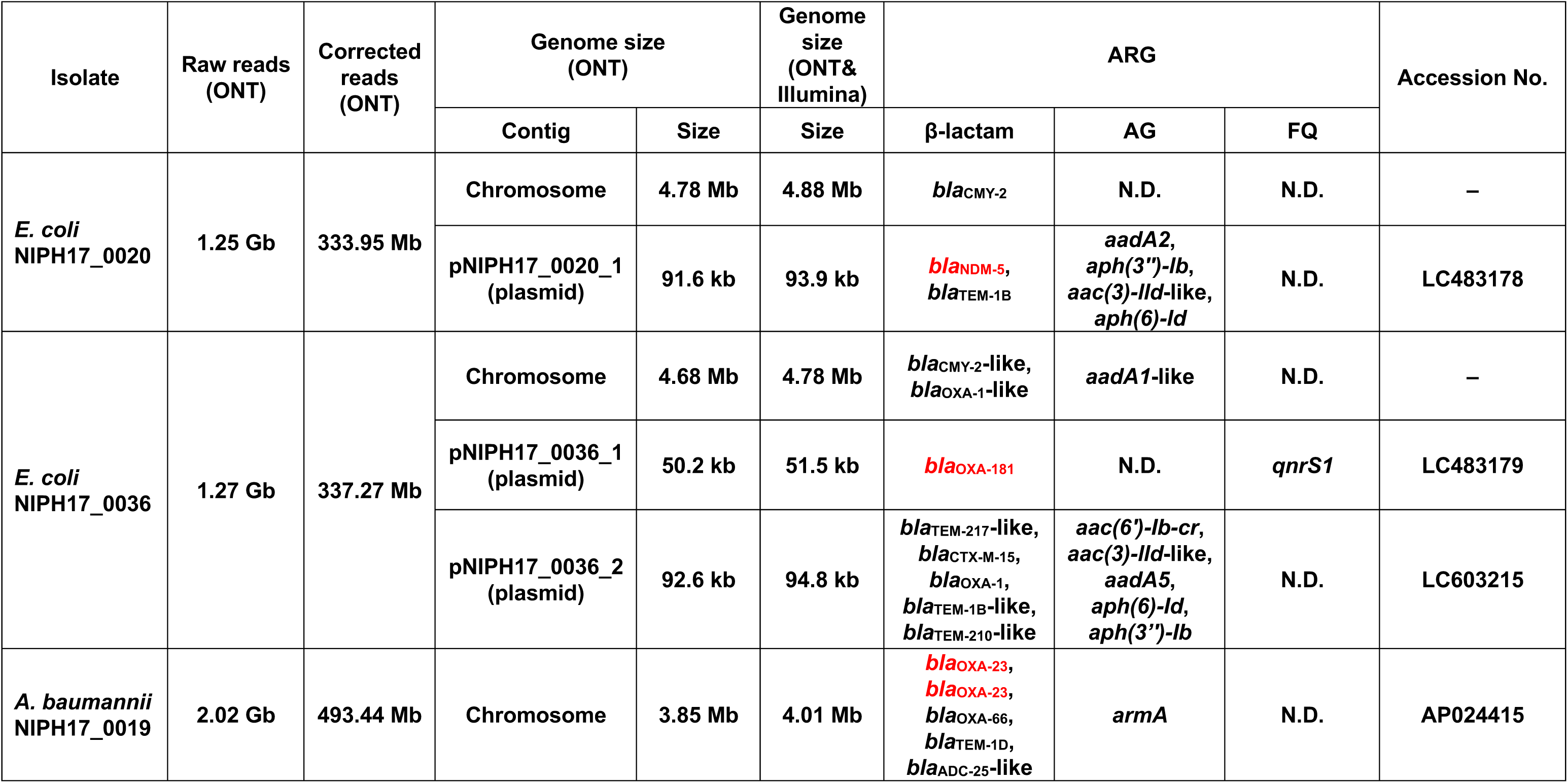
Summary of whole-genome sequencing and bioinformatics analysis of AMR bacteria isolated in Cambodia in this study. Bacterial isolates, nucleotides of raw reads of MinION sequencing and the corrected reads, genome size of the uncorrected sequence (from on-site ONT analysis) and corrected sequence (from subsequent ONT+Illumina analysis) are shown. Also, AMR genes (ARG) associated with resistance to β-lactams, aminoglycosides (AG), and fluoroquinolones (FQ), and accession nos. of sequences are shown.

| Isolate | Raw reads (ONT) | Corrected reads (ONT) | Genome size (ONT) |  | Genome size (ONT& Illumina) | ARG |  |  | Accession No. |
| --- | --- | --- | --- | --- | --- | --- | --- | --- | --- |
| | | | Contig | Size | | β-lactam | AG | FQ | |
| *E. coli* NIPH17_0020 | 1.25 Gb | 333.95 Mb | Chromosome | 4.78 Mb | 4.88 Mb | *bla*<sub>CMY-2</sub> | N.D. | N.D. | – |
|  |  |  | pNIPH17_0020_1 (plasmid) | 91.6 kb | 93.9 kb | *bla*<sub>NDM-5</sub>,<br>*bla*<sub>TEM-1B</sub> | *aadA2*,<br>*aph(3'')-Ib*,<br>*aac(3)-IId*-like,<br>*aph(6)-Id* | N.D. | LC483178 |
| *E. coli* NIPH17_0036 | 1.27 Gb | 337.27 Mb | Chromosome | 4.68 Mb | 4.78 Mb | *bla*<sub>CMY-2</sub>-like,<br>*bla*<sub>OXA-1</sub>-like | *aadA1*-like | N.D. | – |
|  |  |  | pNIPH17_0036_1 (plasmid) | 50.2 kb | 51.5 kb | *bla*<sub>OXA-181</sub> | N.D. | *qnrS1* | LC483179 |
|  |  |  | pNIPH17_0036_2 (plasmid) | 92.6 kb | 94.8 kb | *bla*<sub>TEM-217</sub>-like,<br>*bla*<sub>CTX-M-15</sub>,<br>*bla*<sub>OXA-1</sub>,<br>*bla*<sub>TEM-1B</sub>-like,<br>*bla*<sub>TEM-210</sub>-like | *aac(6')-Ib-cr*,<br>*aac(3)-IId*-like,<br>*aadA5*,<br>*aph(6)-Id*,<br>*aph(3'')-Ib* | N.D. | LC603215 |
| *A. baumannii* NIPH17_0019 | 2.02 Gb | 493.44 Mb | Chromosome | 3.85 Mb | 4.01 Mb | *bla*<sub>OXA-23</sub>,<br>*bla*<sub>OXA-23</sub>,<br>*bla*<sub>OXA-66</sub>,<br>*bla*<sub>TEM-1D</sub>,<br>*bla*<sub>ADC-25</sub>-like | *armA* | N.D. | AP024415 |

**Figure S1.**
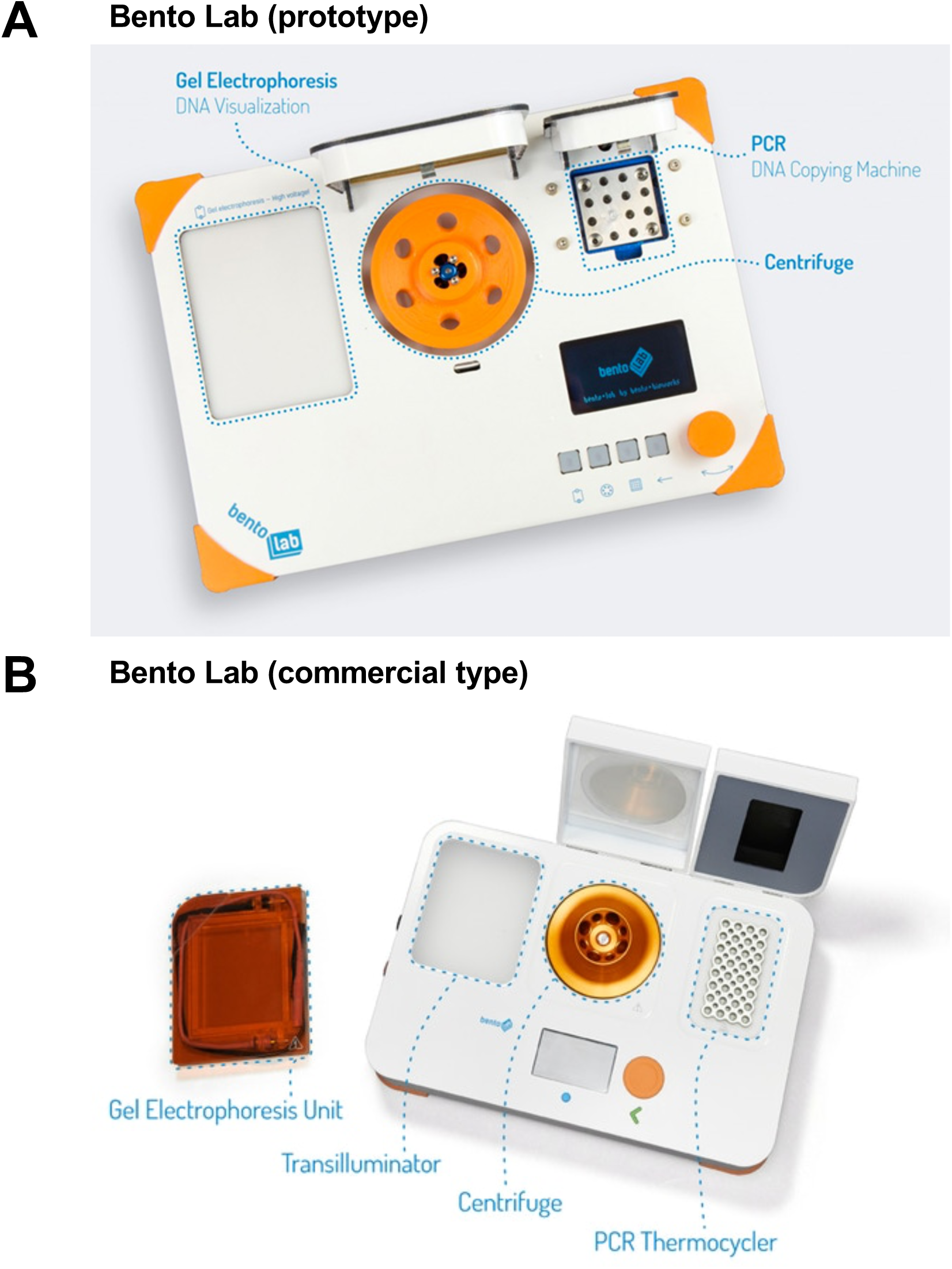
Bento Lab for on-site genomic epidemiological analysis of AMR bacteria in Cambodia. (A) The prototype used in this study and (B) current commercial type are shown.

**Figure S2.**
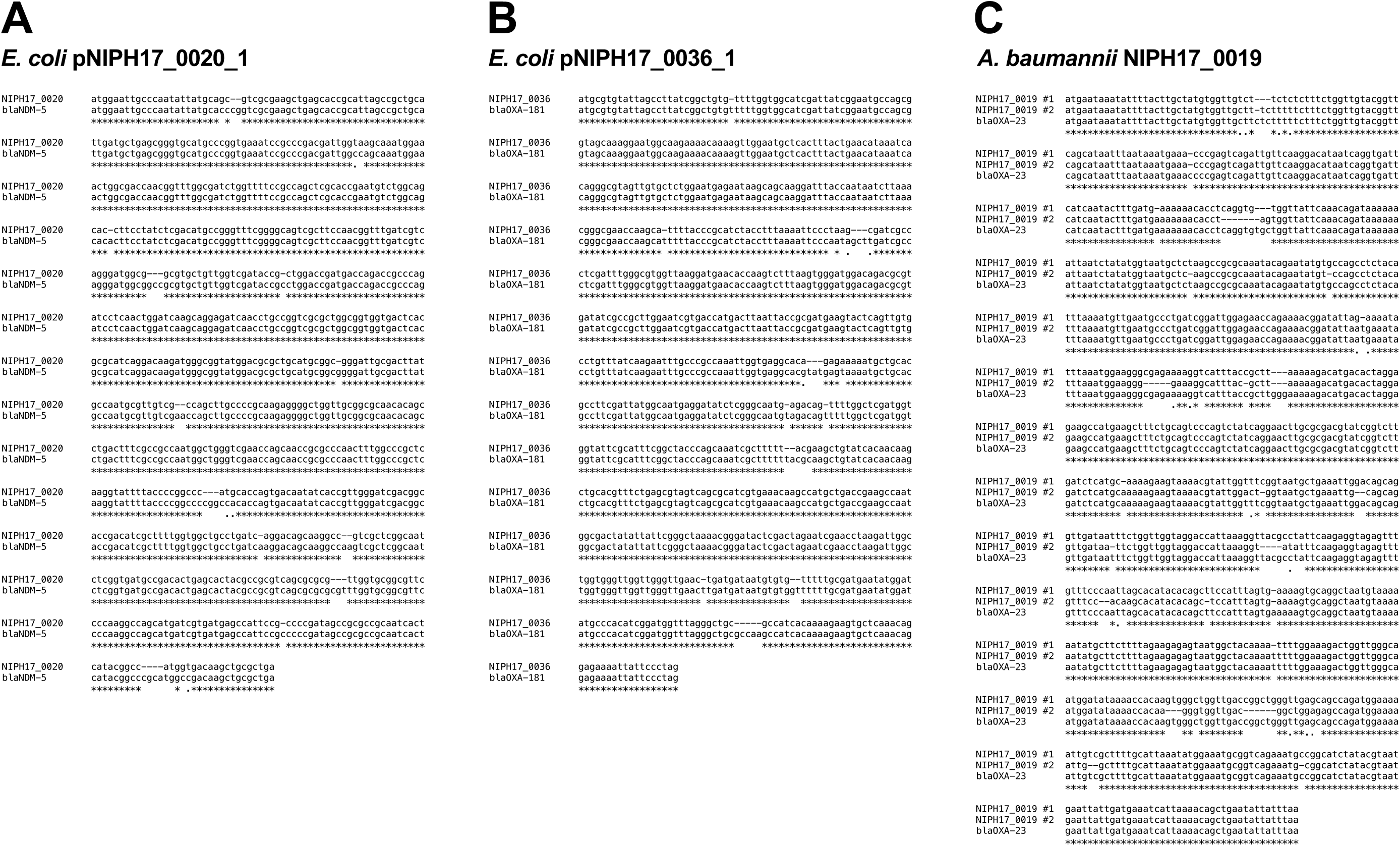
Multiple sequence alignment of carbapenemase genes analyzed by MAFFT v7.475. (A) Comparison between the *bla*_NDM-5_-like gene in *E. coli* pNIPH17_0020_1 (from on-site ONT analysis) and the reference gene (*bla*_NDM-5_ in accession no. JN104597), (B) comparison between the *bla*_OXA-181_-like gene in *E. coli* pNIPH17_0036_1 (from on-site ONT analysis) and the reference sequence (*bla*_OXA-181_ in accession no. CM004561), and (C) comparison between the *bla*_OXA-23_-like sequence in *A. baumannii* NIPH17_0019 (from on-site ONT analysis) and the reference sequence (*bla*_OXA-23_ in accession no. AY795964) are shown.

**Figure S3.**
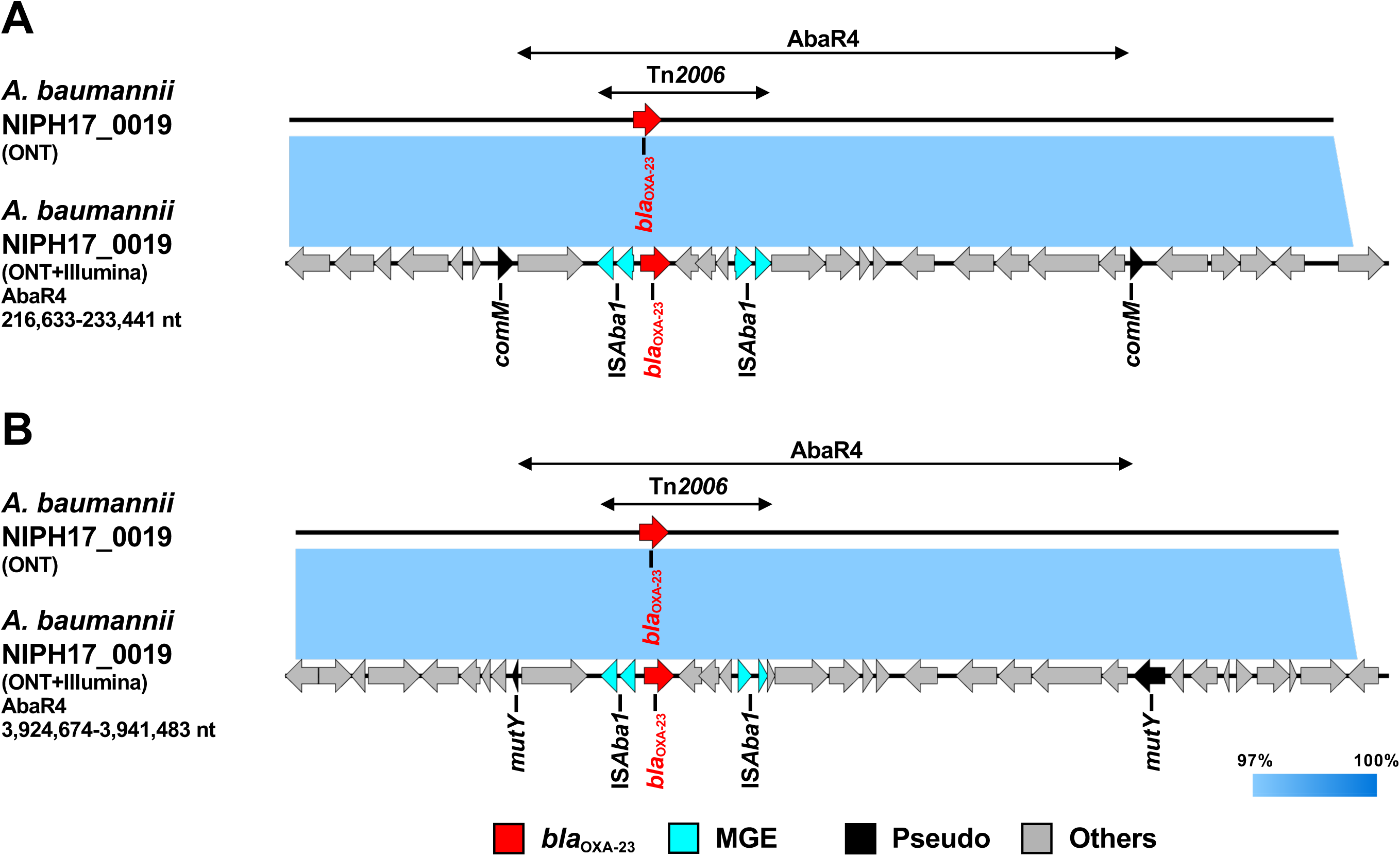
Linear comparison of AbaR4-containing genomic regions in *A. baumannii* NIPH17_0019 harboring two copies of *bla*_OXA-23_. Two sets of AbaR4 with *bla*_OXA-23_ in *A. baumannii* NIPH17_0019: (A) 216,633-233,441 nt region inserted within *comM* and (B) 3,924,674-3,941,483 nt region inserted within *mutY* in accession no. AP024415, and (A and B) comparison of both the uncorrected sequences (from on-site ONT analysis) and corrected sequences (from subsequent ONT+Illumina analysis) are shown. Red, blue, black, and gray arrows indicate carbapenemase gene (*bla*_OXA-23_), mobile gene elements (MGE), pseudogenes disrupted by AbaR4 insertion (Pseudo), and other genes (Others), respectively. The colors in comparison of sequences show percent identity.

